# EEG denoising during transcutaneous auricular vagus nerve stimulation across simulated, phantom and human data

**DOI:** 10.1101/2024.05.13.593884

**Authors:** Joshua P. Woller, David Menrath, Alireza Gharabaghi

**Affiliations:** Institute for Neuromodulation and Neurotechnology, University Hospital Tübingen (UKT), Faculty of Medicine, University Tübingen, 72076 Germany; Max-Planck-Institute for Biological Cybernetics, 72076 Tübingen, Germany; Center for Bionic Intelligence Tübingen Stuttgart (BITS), 72076 Tübingen, Germany; German Center for Mental Health (DZPG), 72076 Tübingen, Germany

**Keywords:** electroencephalography (EEG), transcutaneous electrical stimulation, auricular vagus nerve stimulation (taVNS), data preprocessing, artifact denoising techniques, stimulation artifact

## Abstract

**Objective:** The acquisition of electroencephalogram (EEG) data during neurostimulation, particularly concurrent transcutaneous electrical stimulation of the auricular vagus nerve, introduces unique challenges for data preprocessing and analysis due to the presence of significant stimulation artifacts. This study evaluates various denoising techniques to address these challenges effectively.

**Methods:** A variety of denoising techniques were investigated, including interpolation methods, spectral filtering, and spatial filtering techniques. The techniques evaluated included low-pass and notch filtering, spectrum interpolation, average artifact subtraction, the Zapline algorithm, and advanced methods such as independent component analysis (ICA), signal-space projection (SSP), and generalized eigendecomposition with stimulation artifact source separation (GED/SASS). The efficacy of these algorithms was evaluated across three distinct datasets: simulated data, data from a gelatin phantom model, and real human subject data.

**Results:** Our findings indicate that GED (SASS) and SSP significantly outperformed other methods in reducing artifacts while preserving the integrity of the EEG signal. ICA and Zapline were effective too, but came with important limitations. These methods demonstrated robustness across different data types and conditions, providing effective artifact mitigation with minimal disruption to other essential signal components.

**Conclusion:** This comprehensive analysis demonstrates the efficacy of advanced spatial filtering techniques in the preprocessing of EEG data during auricular vagus nerve stimulation. These methods offer promising avenues for enhancing the quality and reliability of neurostimulation-associated EEG data, facilitating a deeper understanding and wider applications in clinical and research settings.

## Introduction

Transcutaneous auricular vagus nerve stimulation (taVNS) is an emerging neurostimulation technique, with potential applications in a range of psychiatric and neurological disorders such as depression and epilepsy (Badran & Austelle, 2022; Farmer et al., 2021; Yap et al., 2020). TaVNS involves the stimulation of a superficial auricular branch of the vagus nerve (ABVN) with short electrical pulses. The ABVN is considered a promising target due to the far-reaching projections and regulation of physiological and cognitive processes that the vagal system offers (Butt et al., 2020). A literature review (Gianlorenco et al., 2022) has evaluated 18 studies of combined EEG and taVNS. Most reviewed studies used a crossover design, and did not analyze EEG data recorded during task-concurrent stimulation. Yet, recent developments in the field of neurostimulation (Deer & Mali, 2016; Denison & Morrell, 2022) aim to transition from pre-configured stimulation protocols to adaptive stimulation protocols, with stimulation tailored to either the current physiological state or specific task phases.

It is important to note that EEG recorded during electrical stimulation may be severly affected by noise (Iturrate et al., 2018), which must be removed prior to analysis. While artifact reduction methods have been systematically evaluated for other neuromodulation techniques, such as transcranial magnetic stimulation (TMS) (Hernandez-Pavon et al., 2022), this has not yet been done for taVNS.

The stimulation pulses of taVNS are relatively short, with a duration of approximately hundreds of microseconds. However, the recorded artifact of a single stimulation pulse in EEG may have a duration of 3-5 *milli*seconds, depending on the hardware and recording settings (see fig. 3). Similar artifact durations have been reported in MEG as well (Keatch et al., 2023). In the power spectrum, the artifacts are visible as peaks at the stimulation frequency and its higher order harmonics (see fig. 4, 5).

A variety of methods have been proposed for the removal of artifacts in taVNS-EEG/MEG recordings. These include notch filtering (Chen et al., 2023; Lloyd et al., 2023; Sharon et al., 2021), independent component analysis (ICA) (Gurtubay et al., 2023), the Cleanline algorithm (Poppa et al., 2022), linear interpolation (Schuerman et al., 2021), and autoregressive modelling (Keatch et al., 2023). A lack of thoroughly evaluated artifact correction methods is recognized as a current challenge for concurrent taVNS-M/EEG studies (Wienke et al., 2023).

In this study, we evaluate a selection of artifact reduction techniques (see table 1). Projection methods such as signal-space projection (Uusitalo & Ilmoniemi, 1997), ICA (Hyvärinen & Oja, 2000), and generalized eigendecomposition (Haslacher et al., 2021), will be considered alongside lowpass and notch filtering, spectrum interpolation (Leske & Dalal, 2019), and Zapline (de Cheveigné, 2020). Additionally, linear interpolation, mean interpolation and average artifact subtraction are explored. The evaluation is conducted on simulated data, EEG phantom data, and human data.

**Table 1:** Artifact Reduction Methods.

| Method | Method Family | Reference |
| --- | --- | --- |
| Signal-Space Projection | Projection | (Uusitalo & Ilmoniemi, 1997) |
| Stimulation-Artifact-Source<br>Seperation | Projection | (Cohen, 2022; Haslacher et al., 2021) |
| Independent Component Analysis | Projection | (Albera et al., 2012; Hyvärinen & Oja, 2000) |
| Zapline | Spectral & Projection | (de Cheveigné, 2020; Klug & Kloosterman, 2022) |
| Lowpass Filter | Spectral | - |
| Notch Filter | Spectral | - |
| Spectrum Interpolation | Spectral | (Leske & Dalal, 2019) |
| Mean Interpolation | Time-domain interpolation | - |
| Linear Interpolation | Time-domain interpolation | - |
| Average Artifact Subtraction | Template-based | (Kohli & Casson, 2015; Sun & Hinrichs, 2016) |

## Methods

### Denoising Methods

Various artifact removal methods are available that may operate on time domain or frequency domain data. Unless otherwise specified, mne-Python implementations of methods were used.

### Lowpass Filtering

Lowpass filtering is a valid and sufficient strategy for those interested in lower frequency activity (de Cheveigné & Nelken, 2019), especially if precise neural event onsets are not a concern (Vanrullen, 2011; Widmann & Schröger, 2012). For this study, a lowpass filter at 50Hz filter was used. This cutoff frequency was selected to retain as much neural signal as possible.

### Notch Filtering

Another approach to address spectrally well-defined noise is the application of one or multiple notch filters. Notch filters may induce artifacts, such as ringing, when faced with transients and discontinuities in the data (de Cheveigné & Nelken, 2019; Kirac et al., 2016), such as the onset of a stimulation train. Notch filters were implemented at the artifact frequency and its harmonics up to 250 Hz.

### Spectrum Interpolation

Spectrum interpolation was proposed as an approach to remove line noise artifacts from M/EEG data (Leske & Dalal, 2019). It uses a linear interpolation in the frequency domain within a band around specified noise frequencies, which is then transformed back to the time domain, yielding a cleaned signal. Spectrum interpolation has been shown to reliably remove line noise, and is computationally efficient. A custom implementation was used for the present study.

### Zapline

Zapline (using denoising source separation; DSS) is a recent method developed to reduce continuous, spectrally defined noise, such as line noise (de Cheveigné, 2020). It involves spectral and spatial filtering of data. By filtering only a spectrally defined subset of the data, it can preserve both the full spectral band, and the full rank of the data. A Python implementation of Zapline was used (dss.dss_line; meegkit). 10 iterations of the Zapline algorithm were run, as single runs did not provide sufficient noise removal. Since Zapline only performs the noise reduction on a spectrally selected subset of the data, repeated iterations was considered acceptable.

### Independent Component Analysis

Independent component analysis (ICA) is a multivariate method that decomposes a measured signal into statistically independent components (Shlens, 2014a) and is often used to clean EEG data (Jiang et al., 2019). For the present analyses, the fastICA algorithm was used. ICs were visually inspected, and components carrying the stimulation artifact were removed.

### Signal Space Projection (SSP)

Signal Space Projection (SSP) is a multivariate approach to seperate neural signals from noise (Uusitalo & Ilmoniemi, 1997). Based on principal component analysis, it creates a noise subspace, and constructs a projection matrix orthogonal to this noise subspace to remove artifacts. SSP can also remove the true signal if not applied carefully (Haumann et al., 2016), especially when the assumption of orthogonality of signal and noise is violated (Jiang et al., 2019). Additionally, knowledge of the artifact timing can be leveraged to estimate the noise subspace. In the following, SSP based on data epoched around single stimulation artifacts is called ‘SSP (event)’, while SSP based on longer stimulation trains iwill be referred to as ‘SSP (raw)’. SSP is real-time compatible (Sudre et al., 2011).

### Generalized Eigendecomposition / Stimulation Artifact Source Separation

Joint diagonalization of two covariance matrices by solving the generalized eigenvalue problem can maximize a predefined contrast (Cohen, 2022). For two matrices A, B the generalized eigenvalue problem is as follows (Ghojogh et al., 2023):

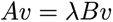

with the generalized eigenvector v, and the generalized eigenvalue λ as solutions. For EEG, A and B could be channel-wise covariance matrices obtained during different conditions. GED provides better separation of correlated sources compared to PCA, as GED can orthogonalize sources that are correlated in channel space (Cohen, 2022)

A GED-based artifact correction method, called stimulation artifact source separation (SASS) has been proposed to remove artifacts from transcranial alternating current stimulation (Haslacher et al., 2021). Haslacher and colleagues performed this GED on band-pass filtered signal covariances from stimulation and non-stimulation periods, and used the resulting GED for artifact removal.

A custom implementation was programmed based on the code of Haslacher et al. (2021). Modifications include the use of a permutation test (n_permutation = 1000, alpha = 0.05) to determine the number of artifact components and the use of covariances computed on broadband data. Covariance matrices were constructed by averaging covariances across artifact-containing and artifact-free non-overlapping epochs around and in between individual stimulation pulses, respectively, each epoch lasting 20ms (GED ‘event’).

### Linear and Mean Interpolation

A basic approach to dealing with artifacts is to replace the affected data segment with a constant or linear trend estimate (Hoffmann et al., 2011). While compatible for some analyses (Zhou et al., 2018), it can reduce the overall variance of the signal and remove the true brain signal. Linear and mean interpolation were implemented using custom Python functions, using a 5ms window around the artifact, which at 25 Hz means 12.5% of the data were imputed.

### Average Artifact Subtraction

Average artifact subtraction (AAS) or template subtraction (TS) refers to an artifact removal method that estimates a template of the artifact, and then subtracts this from the signal. It is often used for concurrent fMRI-EEG to remove the gradient artifacts from EEG (Kraljič et al., 2022), but can also be used for stimulation artifacts, e.g. in tACS (Helfrich et al., 2014), DBS (Sun & Hinrichs, 2016), or TMS (Vernet & Thut, 2014). The subtraction template is constructed by averaging artifact-containing epochs. Hence, AAS works best when the artifact shape is stable over time. Residual artifacts can arise if assumptions are violated (Steyrl & Müller-Putz, 2019), e.g., due to misalignment of template and artifact or changes in artifact shape. For artifacts that contain sharp transients, like electrical stimulation artifacts, misalignment can reintroduce artifacts into the data. For electrical stimulation artifacts like taVNS, the assumption of temporal stability of the measured artifact shape can be systematically violated, as waveform undersampling, resampling and antialiasing filters can modulate the artifact shape (Vernet & Thut, 2014). The template can either be estimated by averaging over all such epochs, or a moving average template can be used. The latter allows for adaptation to slow changes in artifact shape, but leads to noisier template estimates.A modified AAS to prevent aliasing of entrained neural oscillatory activity was used (see appendix C for details). A sliding window of 20 stimulation events was used to estimate the artifact template.

## Data

### EEG Simulation

Data were simulated using custom scripts and mne-Python. Neural sources in (1) the right frontal pole (FPr) and (2) the rostral anterior cingulate cortex (rACC) were selected. 30 oscillatory beta events (25Hz) of 1 second duration were generated at the sources and projected to the sensor level. Correlated noise was added. Data were simulated for 60 EEG channels at a sampling rate of 10kHz. A 5s on 5s off stimulation, 25Hz regime was simulated, and the resulting time course was added to the EEG data. Data were simulated once with and once without the ringing artifacts resulting from the stimulation for each source (see appendix B for details).

### Phantom

To evaluate the performance of the different artifact correction methods under controlled, but physically more realistic conditions, a gelatin head phantom was constructed, according to published instructions (Hairston & Yu, 2019).

The phantom was cast using ballistic gelatine (Gelatine Ballistic 3, Kremer Pigmente GmbH, Aichstetten, Germany) prepared with salt water (40.5 g NaCl in 4.5 L of distilled water).

In a passive configuration, a head-shaped inlet printed from ABS was added to the mold, to better approximate the spread of electrical signals on the scalp.

In the active configuration, six auxiliary cables were embedded in the gelatin. An external audio-card was used to inject 10 Hz signals into the phantom during the experiment. 10 Hz oscillations were chosen as the alpha range is of interest in recent taVNS research (Lloyd et al., 2023; Sharon et al., 2021). To achieve realistic impedance levels, the phantom was powdered (Penaten powder, containing corn starch, aloe extract and tricalcium phosphate) and a layer of tubular bandages (Stülpa head bandages, Paul Hartmann GmbH, Germany) was applied. This provided to a stable impedance range of 3-10 kOhm and resulted in clearly visible stimulation artifacts in the EEG.

The taVNS stimulator was attached to the phantom ear and held in place with bandages. Stimulation was applied at 1s on, 1s off intervals, at 25 Hz with 3mA (see table 2). An actiCHamp amplifier with a 64 channel Acticap slim EEG system (Brain Products GmbH, Inning, Germany) was used for the phantom experiment. Data were recorded at a sampling rate of 25 kHz.

**Table 2:** Stimulation Parameters.

| Subject | Current | Pulse Width | Train Duration | Frequency |
| --- | --- | --- | --- | --- |
| Human | 1.2mA | 250µs | 5s on, 5s off | 25Hz |
| Phantom | 3mA | 250µs | 1s on, 1s off | 25Hz |

### Human

The participant met the inclusion criteria and gave informed consent to take part in the measurements. Data were collected as part of a study approved by the Ethics Committee of the University Hospital Tübingen (178/2023BO1).

The stimulation intensity was adjusted to produce a noticeable, but not painful sensation. The subject was asked to focus on a fixation cross presented on a screen. After a short baseline EEG measurement (∼1 min), stimulation was started. Data were recorded using the BrainAmp amplifier (5 kHz, 1 kHz high cut-off) and a 32 channel Acticap EEG system (Brain Products GmbH, Inning, Germany). Electrodes were placed according to the 10-20 system. Three such data segments (at least 5 min duration) were recorded. Stimulation was applied in intervals of 5 s on, 5 s off at 25 Hz with 1.2 mA (see table 2).

### taVNS Stimulation

For the application of taVNS, the tVNS-R device (tVNS technologies GmbH, Erlangen, Germany) was used. The device generates biphasic stimulation pulses at a specified frequency (1-40Hz), with a minimum duration of of 1s per stimulus train. Between stimulation phases, the device applies a small electrical current every second to measure the impedance, and adjust the voltage accordingly. The stimulation current is delivered to the ear using an earpiece (see fig. 1) with two titanium ball electrodes.

**Figure 1:**
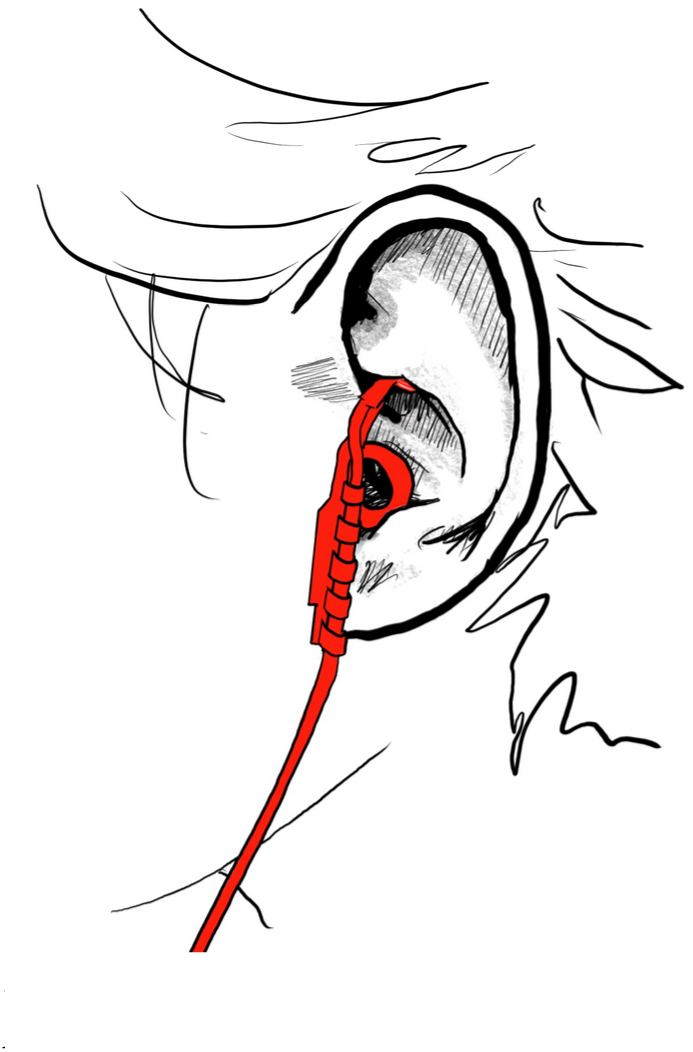
Application of the classical ear piece to the cymba conchae. Illustration by Tristan Wippermann, used with permission.

Electrodes are placed at the cymba conchae (see fig. 1) for stimulation. The tVNS-R stimulator was connected to the experimenter’s computer via Bluetooth using the manufacturer’s software (tvnsmanager.exe, tVNS technologies GmbH, Erlangen, Germany).

### Data processing

Data were processed in Python using the mne-Python, meegkit, numpy and scipy libraries, and custom functions.

Data were cropped to a maximum duration of 10 minutes and resampled to 5kHz (anti-aliasing filter at 2.5kHz, high-pass filter at 1Hz). Power line noise was removed via spectral interpolation at 50Hz, with a bandwidth of 1Hz (Leske & Dalal, 2019). No re-referencing was applied.

Since the tVNS-R device does not provide a trigger channel, stimulation pulse timings had to be inferred from the data. A custom PCA-based thresholding procedure was developed (see appendix A), which annotated the data with stimulation and impedance pulse timings .

## Statistical Evaluation

### General considerations

To evaluate denoising procedures, cleaned data were compared to baseline or ground truth data. Suitable artifact reduction techniques should show a small difference between the cleaned and baseline spectra. The average signal in the cleaned artifact epochs should be low. Furthermore, oscillatory activity overlapping with the stimulation frequency should not be affected. Additionally, denoising methods should be conceptually scalable to different stimulation frequencies, such as high-frequency bursts (Sclocco et al., 2020).

### Comparison of Power Spectral Density (PSD)

PSDs were calculated for the frequency range of 1 Hz – 250 Hz using Welch’s method. Similarity between spectra was characterized by the root mean squared error (RMSE) as a measure of total difference and the Earth Movers Distance (EMD, or Wasserstein metric) as a measure of spectral shape similarity (Majewski et al., 2018).

### Power at specific frequencies of interest

In addition, the difference in amplitude between the spectra at specific frequencies, namely at 10 Hz as a representative of the alpha range (Lloyd et al., 2023; Sharon et al., 2021), and at the stimulation frequency of 25 Hz and its harmonics up to 250 Hz is assessed. In the time domain, the absolute mean signal across stimulation epochs is compared before and after cleaning, with a significant reduction in mean amplitude expected for successful artifact removal.

### Visual inspection

Furthermore, spectra and sections of the EEG time course are plotted and visually inspected for residual artifacts and anomalies. Average artifacts extracted from stimulation epochs are inspected. Additionally, time-frequency plots of the average EEG field for different denoising strategies are presented.

## Results

All methods were able to reduce artifacts. However, when comparing their spectra to the baseline, clear differences in the accuracy of the different methods emerged (see table 3, and fig. 2). GED (SASS) produced the highest similarity to the baseline spectrum and the lowest RMSE. SSP was also highly accurate in reconstructing spectral features, with an advantage when using epoched data (‘SSP (event)’, EMD = 0.57, sd = 0.66, RMSE = 1, sd = 0.788) compared to SSP performed on a continuous segment (‘SSP (raw), EMD = 0.646, sd = 0.826, RMSE = 1.246, sd = 1.089). In the human data, GED achieved almost complete recovery of the baseline spectrum (see fig.4). Compared to SSP, ICA provided slightly better performance (EMD = 0.558, sd = 0.805, RMSE = 0.795, sd = 1.003). This came at the cost of a greater loss of rank for some datasets, as more components were discarded for ICA than for SSP, especially for human data (see table6). Both Zapline (EMD = 2.479, sd = 1.578, RMSE = 2.942, sd = 1.759) and notch filtering (EMD = 1.907, sd = 0.646, RMSE = 5.29, sd = 0.445) failed to preserve spectral features and showed large dissimilarity values. This was expected for notch filtering. Visual inspection of the Zapline data cleaned shows a loss of spectral power restricted to frequencies above 50 Hz (see fig.5).

**Figure 2:**
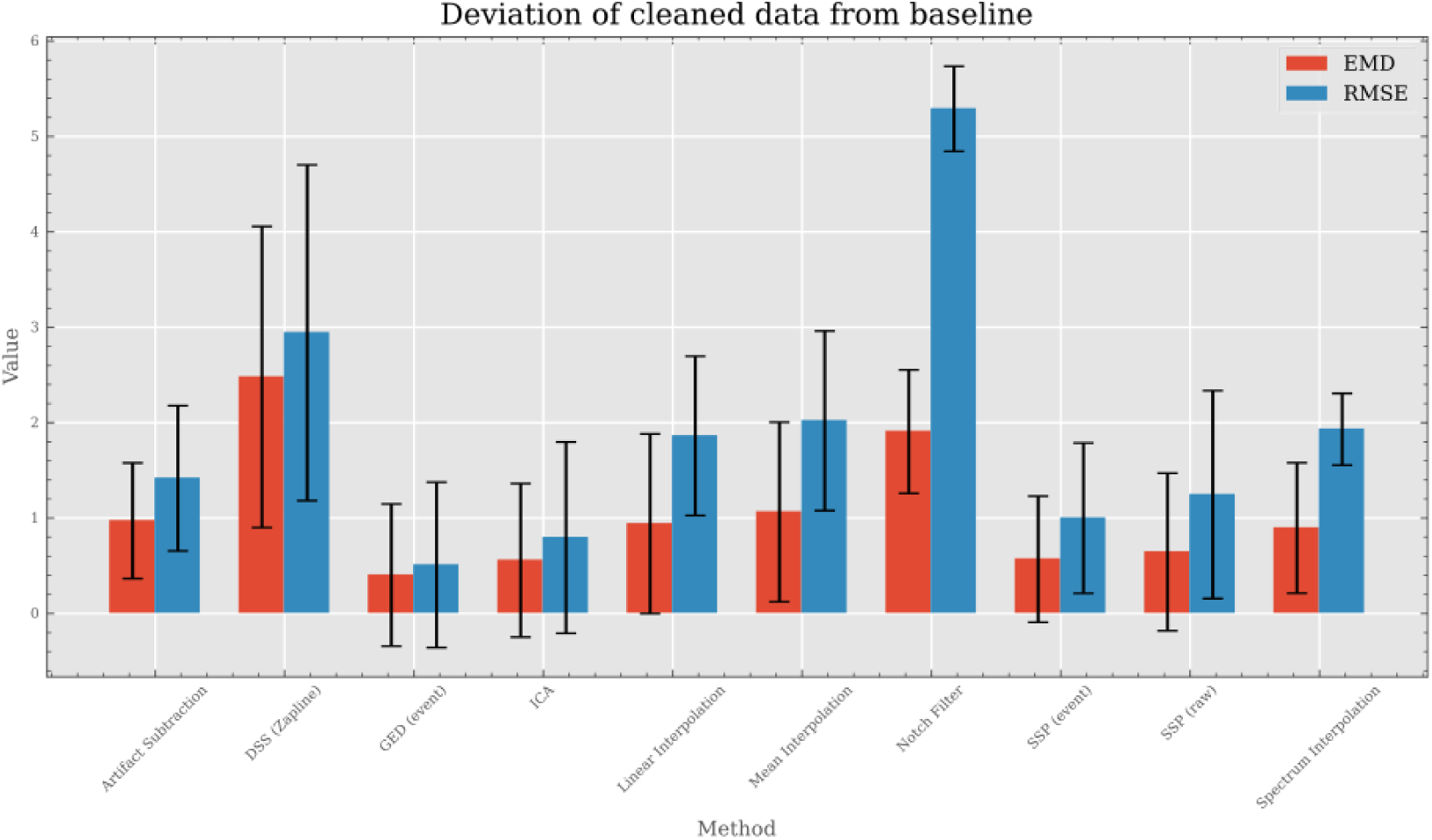
Average PSD Dissimilarity values across all datasets for different artifact reduction methods. Lowpass filtering was omitted from this comparison.

**Figure 3:**
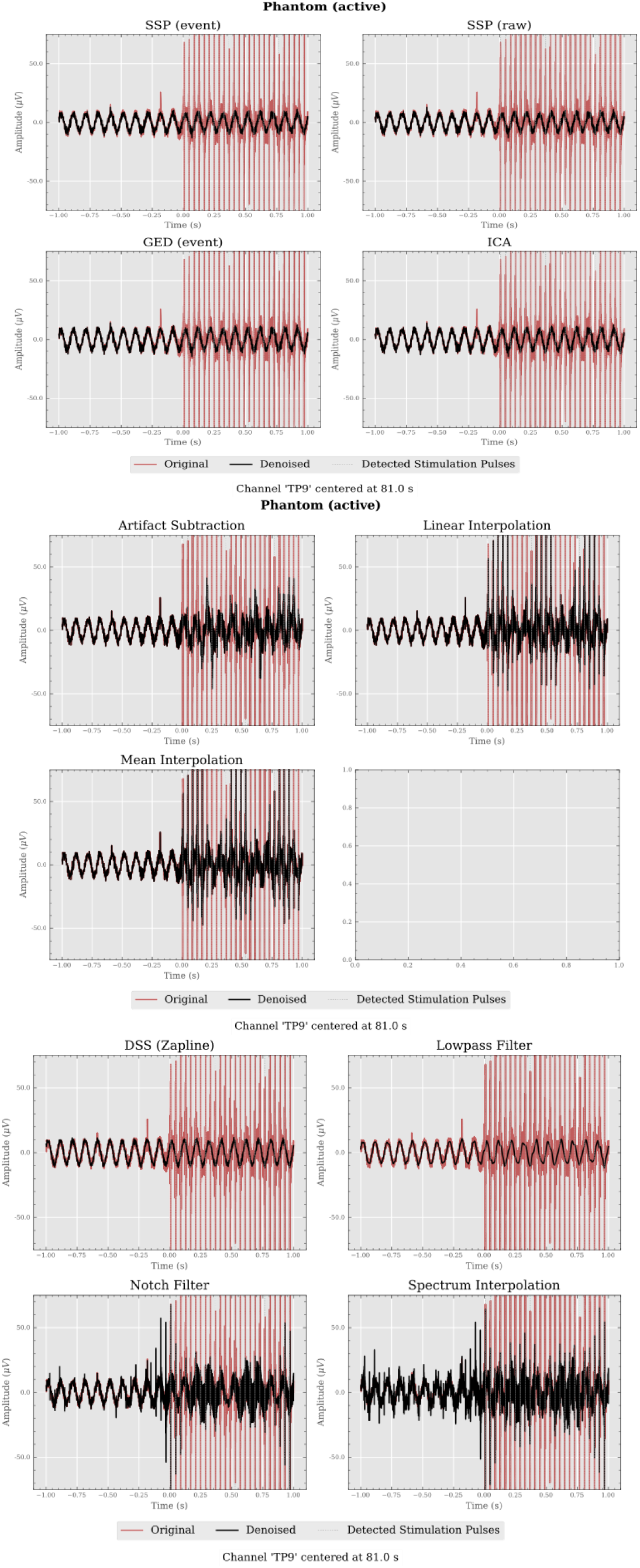
2s samples of an active phantom with 10Hz oscillations. Original signal is shown in red, the cleaned signal in black. (Upper) Projection methods. (Middle) Interpolation and AAS. (Lower) Spectral Methods.

**Figure 4:**
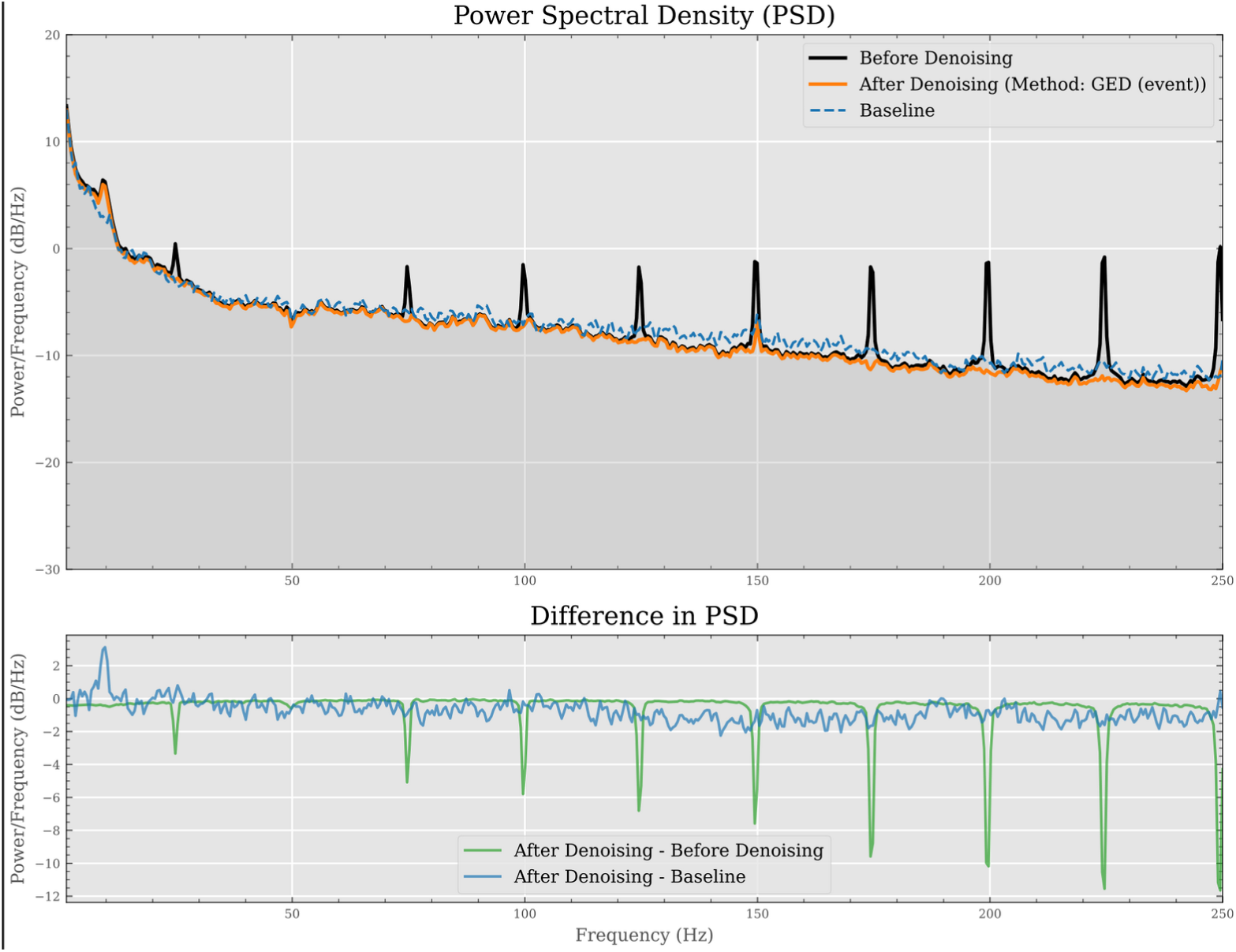
Comparison of spectra before and after denoising with GED (human).

**Figure 5:**
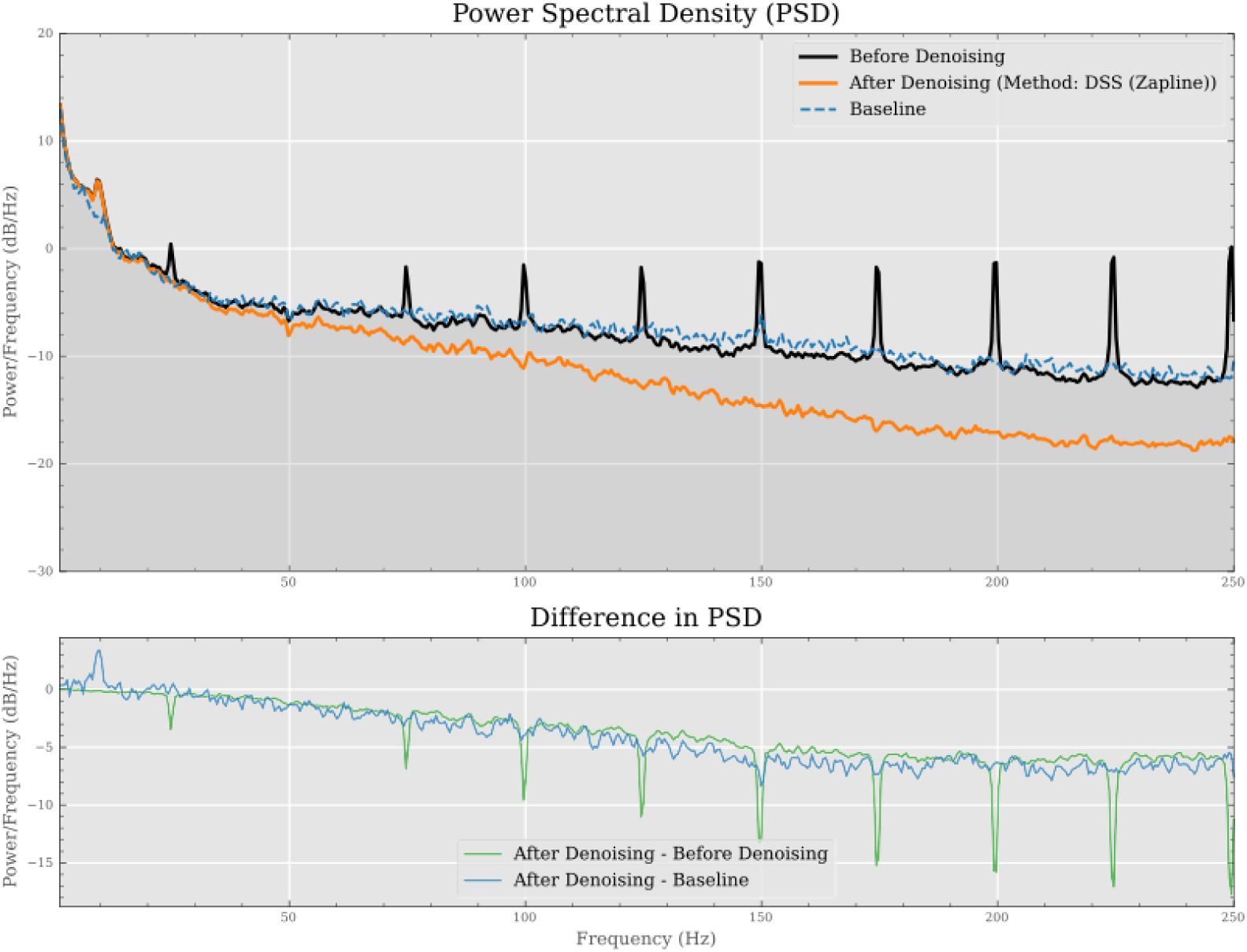
Comparison of spectra before and after denoising with Zapline (human).

**Table 3:** Dissimilarities of cleaned vs baseline spectra across all subjects. Smaller values indicate greater similarity. Smaller standard errors indicate less variance across subjects.

| Method | EMD |  | RMSE |  |
| --- | --- | --- | --- | --- |
|  | mean | sd | mean | sd |
| AAS | 0.97 | 0.61 | 1.42 | 0.76 |
| Zapline | 2.48 | 1.58 | 2.94 | 1.76 |
| GED (event) | 0.4 | 0.75 | 0.51 | 0.87 |
| ICA | 0.56 | 0.81 | 0.8 | 1 |
| Linear Interpolation | 0.94 | 0.94 | 1.86 | 0.84 |
| Mean Interpolation | 1.06 | 0.94 | 2.02 | 0.94 |
| Notch Filter | 1.91 | 0.65 | 5.29 | 0.45 |
| SSP (event) | 0.57 | 0.66 | 1 | 0.79 |
| SSP (raw) | 0.65 | 0.83 | 1.25 | 1.09 |
| Spectrum Interpolation | 0.9 | 0.68 | 1.93 | 0.38 |

Visual inspection of the time domain signals reveals differences of between the artifact reduction techniques (see fig.3). In the active phantom SSP, ICA, GED and Zapline recovered the underlying 10Hz oscillation during the stimulation period, while linear and mean interpolation failed to remove all peaks of the stimulation artifact. AAS recovered the time course, mixed with residual artifacts. Notch filtering and spectral interpolation yielded strong artifacts at the edges of the stimulation trains.

In the time domain, the absolute values of the signal within stimulation epochs (± 10ms, see fig. 6, 7) were averaged before and after denoising, and the difference was computed. All methods substantially reduced the absolute mean signal (see table 4). The highest reduction was achieved by ICA (6.17 µV), followed by Zapline (6.03 µV) and GED (6.02µV). The mean difference between data at selected frequencies was calculated on PSDs of individual subjects using a 1 Hz bandwidth, and then averaged across subjects (see figure 8; see table 5 for averages across frequencies). Overall SSP, ICA, GED and AAS had low errors across frequencies. Zapline, linear and mean interpolation had errors especially at higher frequencies. Spectrum interpolation had an anomalous reduction at the 25 Hz frequency.

**Figure 6:**
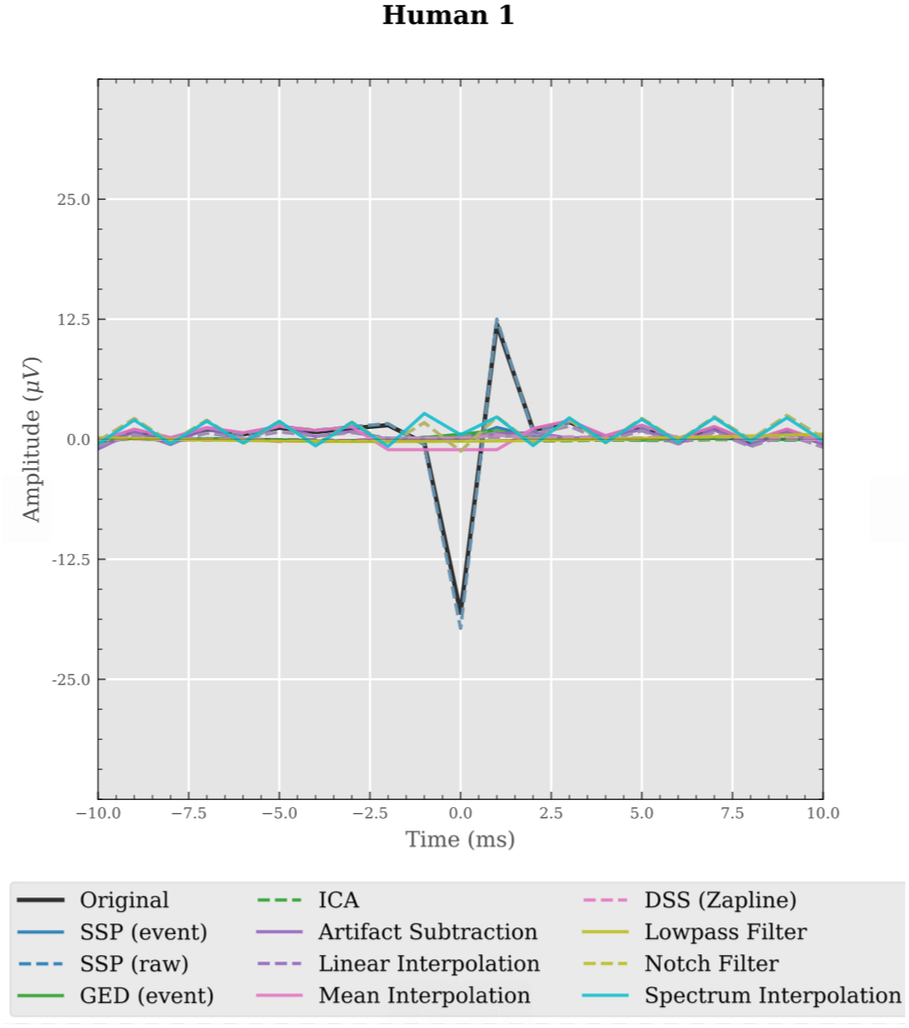
Average Stimulation artifact after various denoising methods (human).

**Figure 7:**
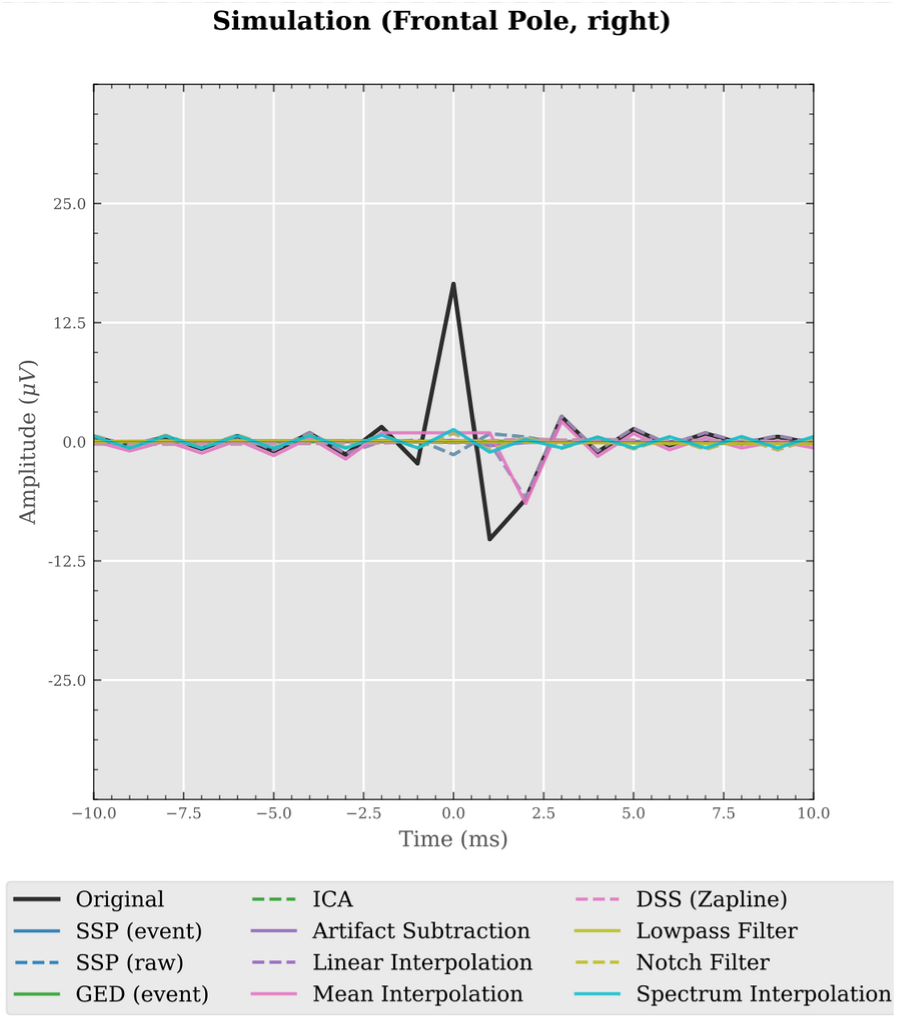
Average Stimulation artifact after various denoising methods (simulation).

**Figure 8:**
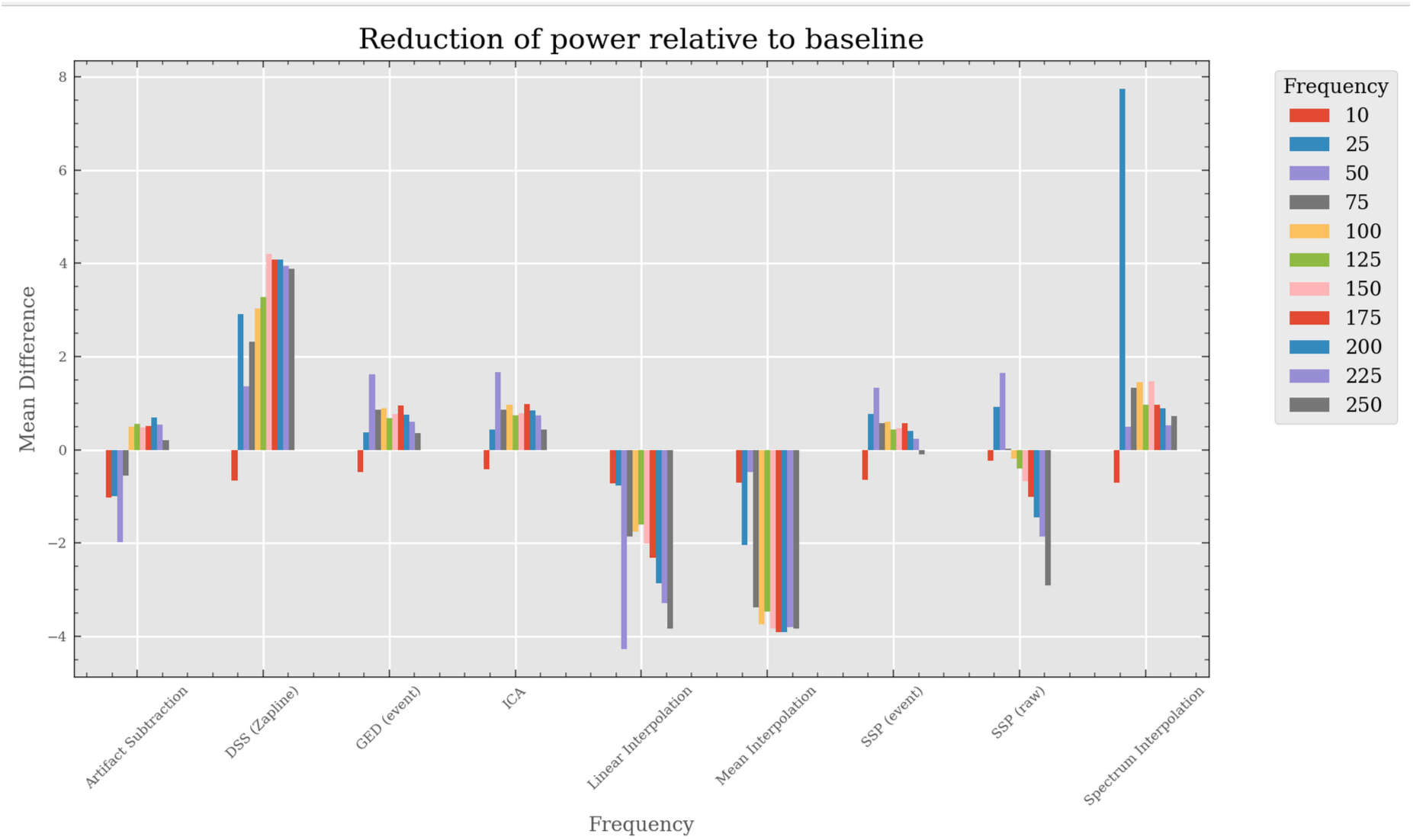
Mean Difference in spectral power relative to baseline, averaged across subjects.

**Table 4:** Average absolute signal during artifact epochs, across all channels for all subjects.

| Method | Absolute mean (μV) | | Difference to original(μV) | |
| --- | --- | --- | --- | --- |
|  | mean | sd | mean | sd |
| AAS | 0.59 | 0.56 | 5.72 | 3.64 |
| Zapline | 0.27 | 0.29 | 6.03 | 3.92 |
| GED (event) | 0.29 | 0.37 | 6.02 | 3.85 |
| ICA | 0.14 | 0.11 | 6.17 | 4.08 |
| Linear Interpolation | 1.8 | 1.03 | 4.51 | 3.23 |
| Lowpass Filter | 0.44 | 0.34 | 5.86 | 3.86 |
| Mean Interpolation | 2.62 | 1.67 | 3.69 | 2.6 |
| Notch Filter | 2.16 | 1.86 | 4.14 | 2.32 |
| SSP (event) | 0.32 | 0.33 | 5.98 | 3.93 |
| SSP (raw) | 2.76 | 4.93 | 3.55 | 3.37 |
| Spectrum Interpol. | 2.33 | 1.93 | 3.98 | 2.26 |

**Table 5:** Mean difference in the frequency bins 10, 25, 50 … 250 Hz (1 Hz bandwidth). Averaged across datasets.

| Method | Mean difference |
| --- | --- |
| Artifact Subtraction | -0.1 |
| DSS (Zapline) | 2.95 |
| GED (event) | 0.67 |
| ICA | 0.73 |
| Linear Interpolation | -2.3 |
| Mean Interpolation | -3.01 |
| SSP (event) | 0.42 |
| SSP (raw) | -0.56 |
| Spectrum Interpolation | 1.44 |

Time-frequency analysis of the simulated data shows that compared to the artifact-containing data (see fig. 9), the eigendecomposition-based methods (see fig. 11 for SSP) and Zapline (fig. 10) in particular preserve ongoing activity in the same frequency range as the stimulation, while successfully removing artifacts. To a lesser extent, this was also successful for AAS (with some residual artifacts, see fig. 13). Spectrum interpolation reduced the power of transient activity overlapping with the stimulation frequency (see fig.12).

**Figure 9:**
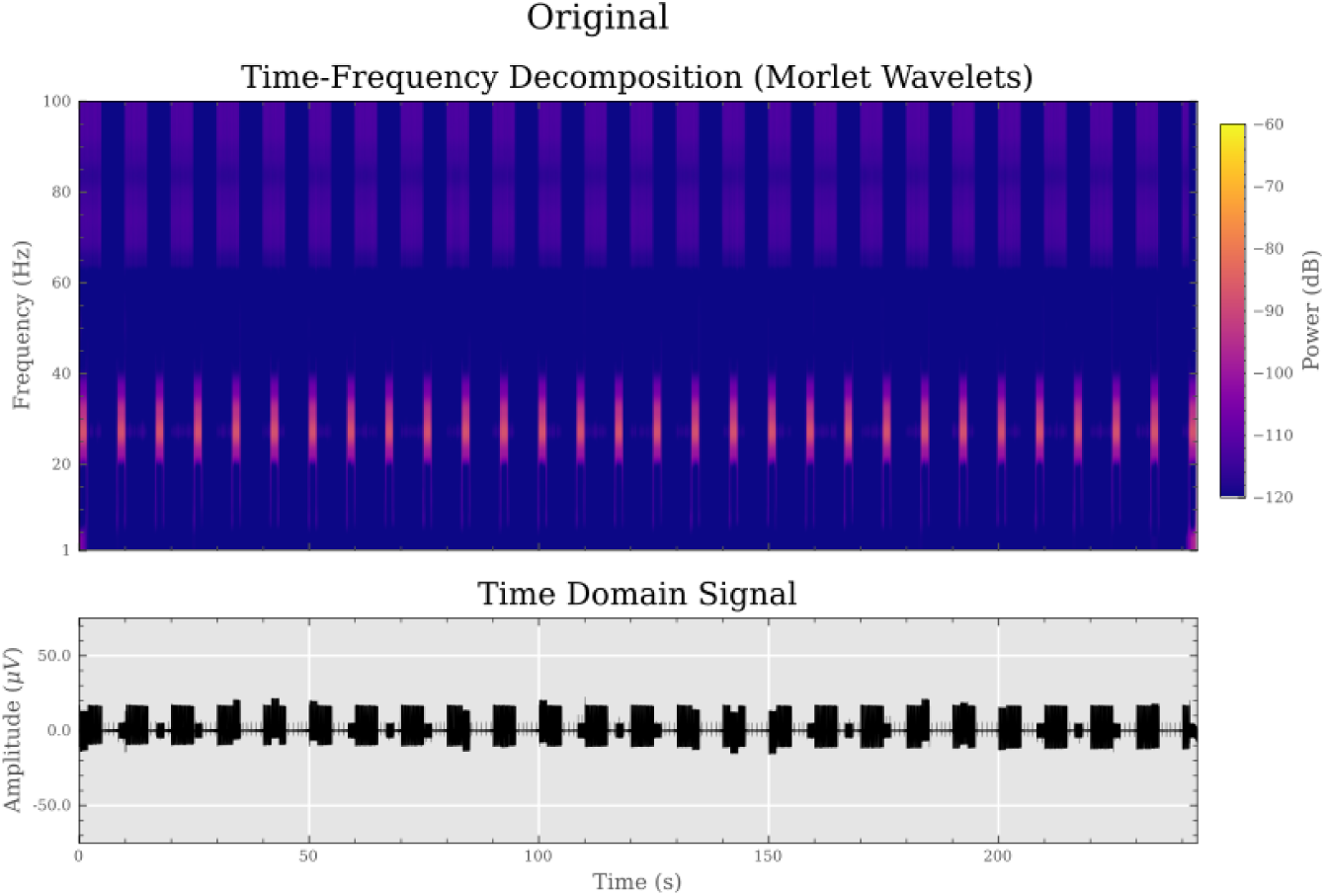
Time frequency decomposition of the original signal averaged across all channels (upper subplot). The average time domain signal is shown below. Transient neural activity of 1s duration at 25Hz was simulated.

**Figure 10:**
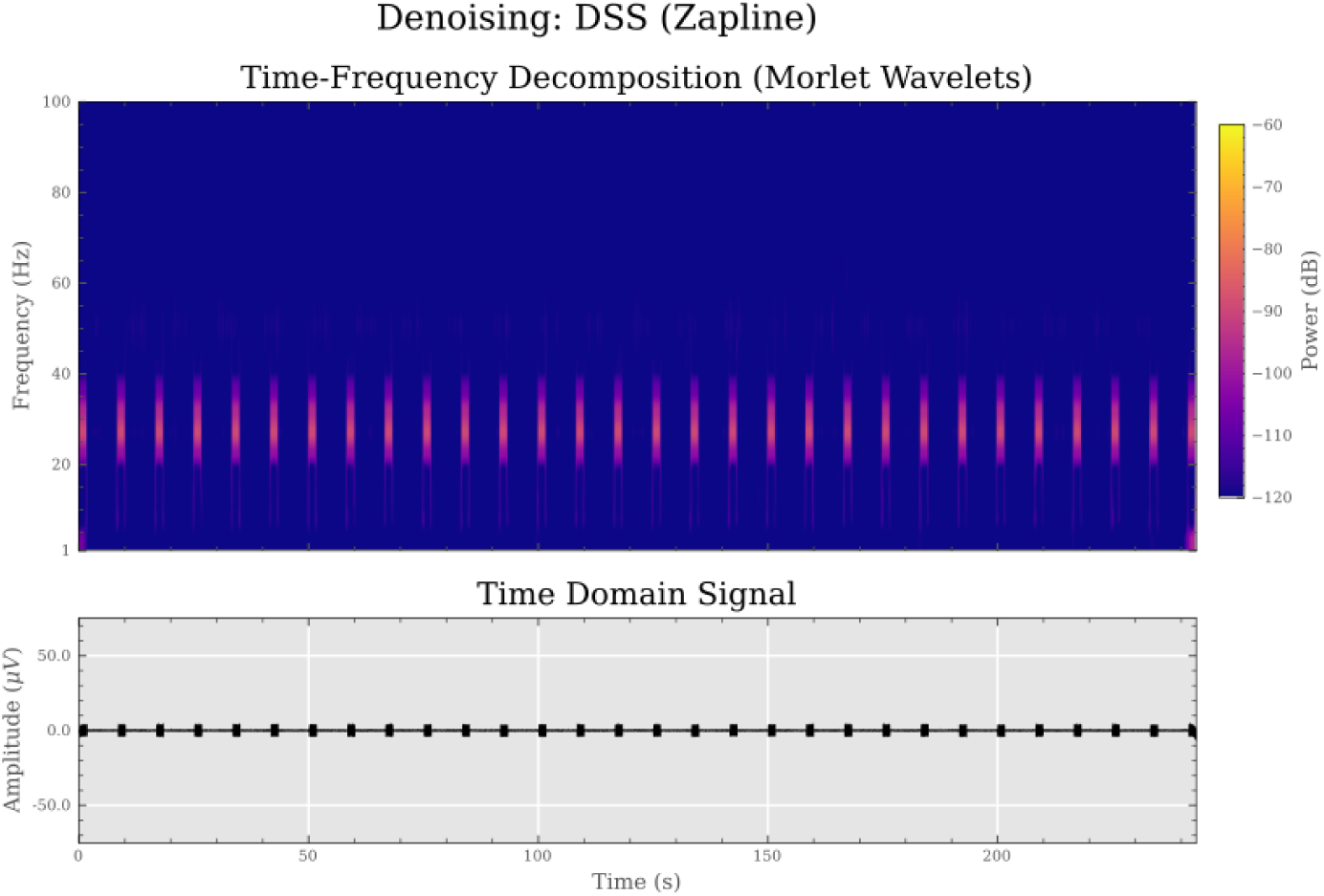
Time frequency decomposition of the signal after Zapline application, averaged across all channels (upper subplot). The average time domain signal is shown below.

**Figure 11:**
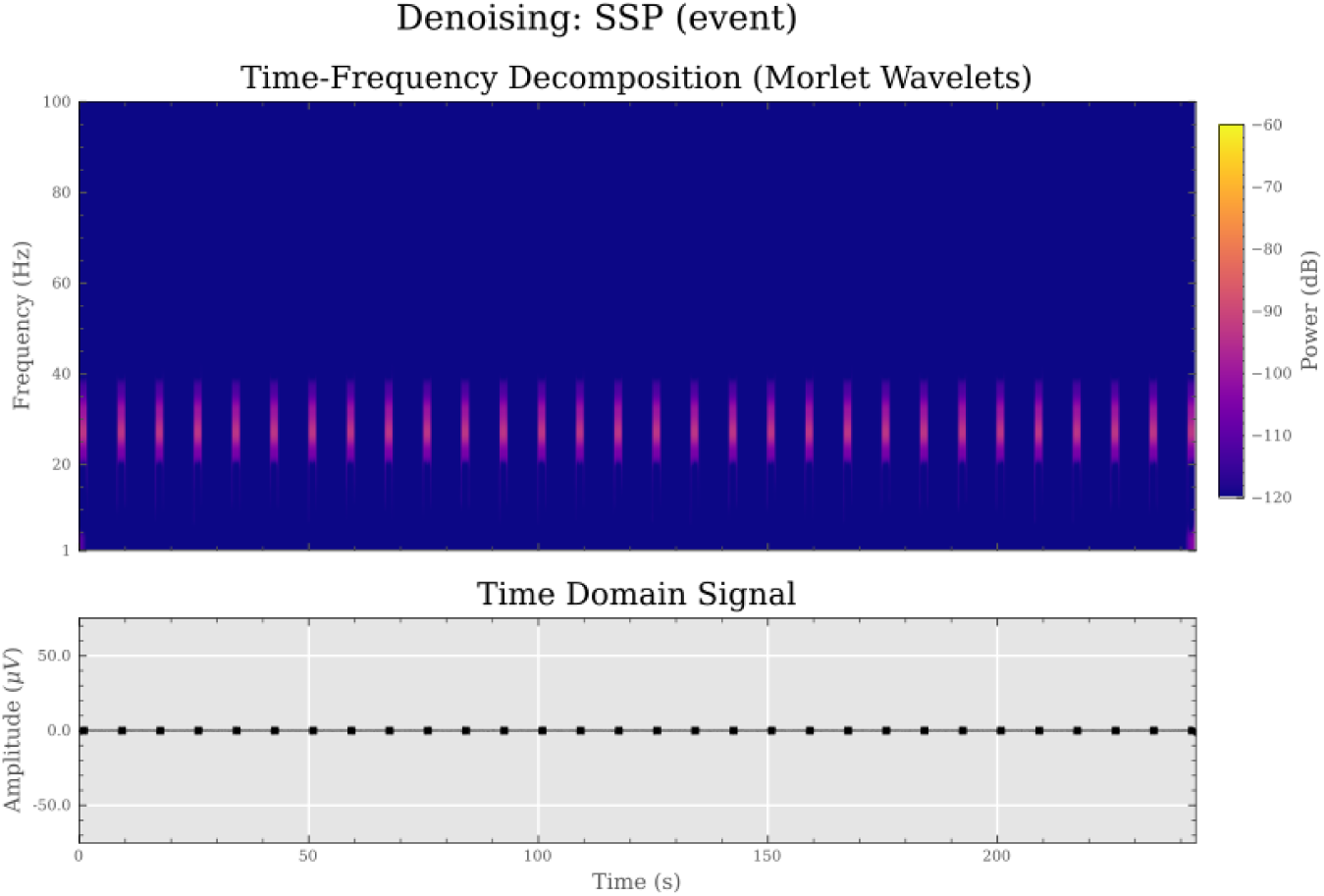
Time frequency decomposition of the signal after cleaning via event-based SSP averaged across all channels (upper subplot). The average time domain signal is shown below.

**Figure 12:**
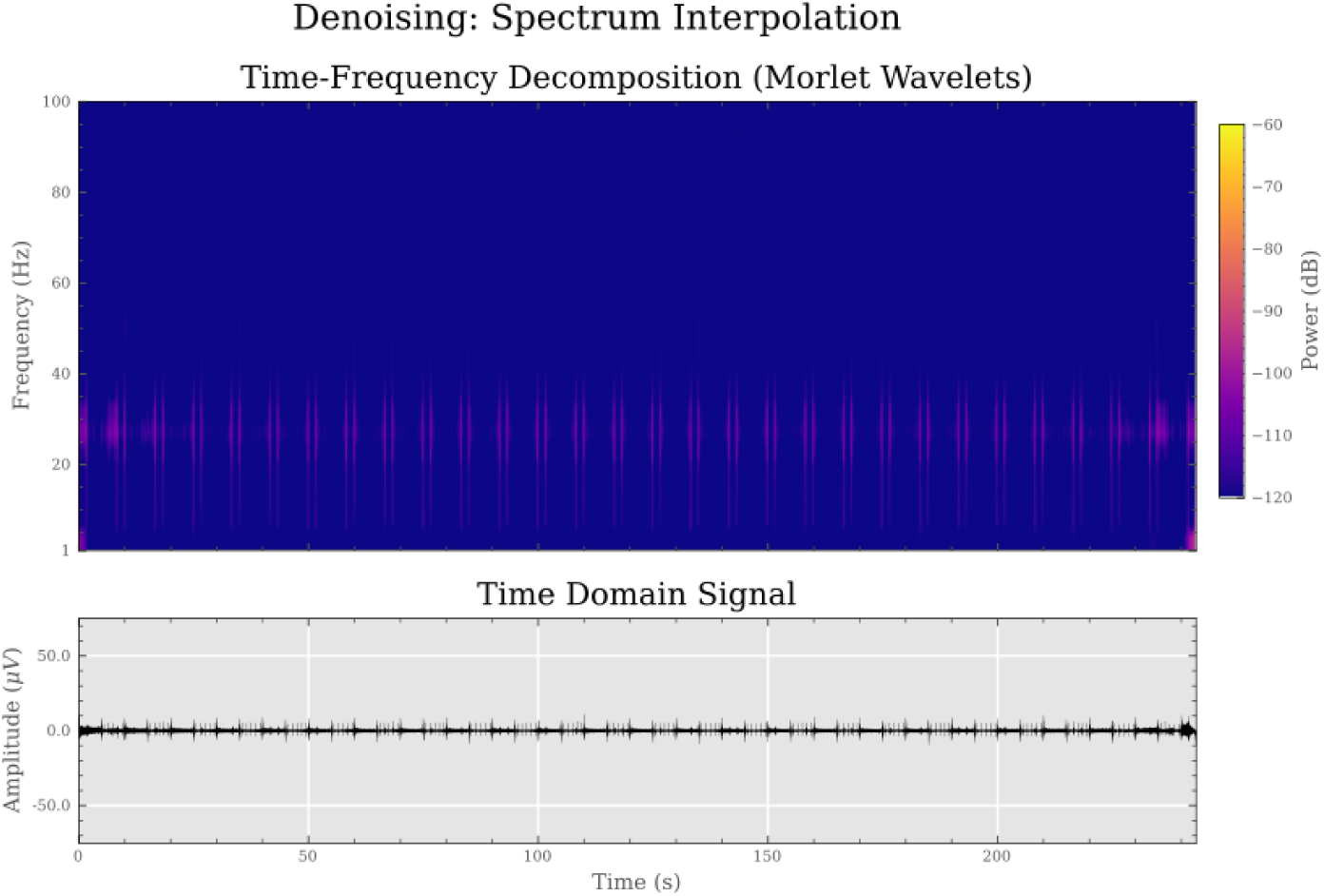
Time frequency decomposition of the signal after spectrum interpolation, averaged across all channels (upper subplot). The average time domain signal is shown below.

**Figure 13:**
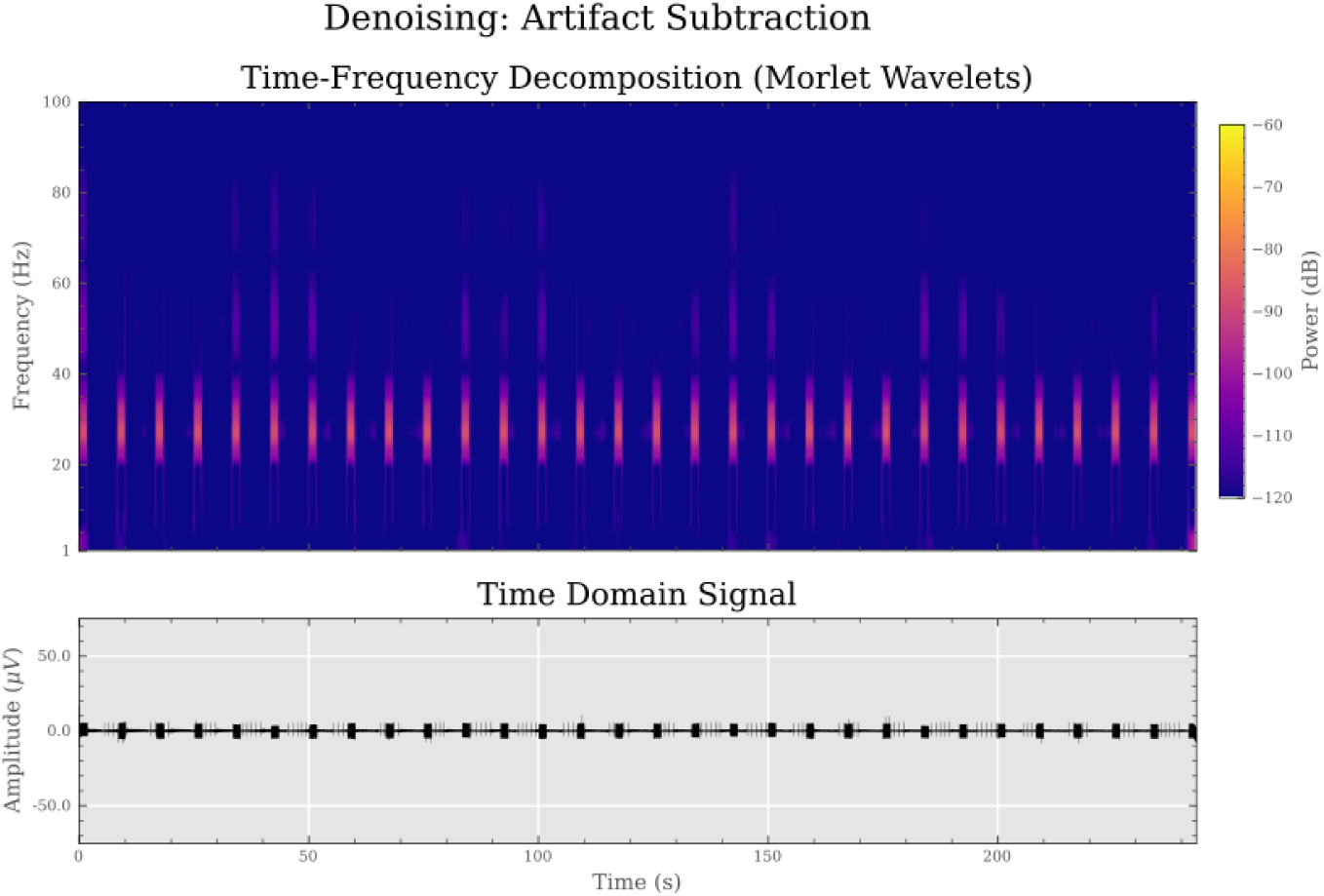
Time frequency decomposition of the signal after average template subtraction, averaged across all channels (upper subplot). The average time domain signal is shown below.

Overall, SSP, GED (SASS) and ICA showed a consistent ability to remove artifacts while preserving the true signal across simulated, phantom and human data.

In SSP and GED, few components needed to be removed, while the number of artifact components was more variable in ICA on real data (see table 6). For SSP and ICA, the removed components were determined by visual inspection, while for GED a permutation approach was used. In the passive phantom, the latter procedure resulted in the removal of 63 components because there was no structured signal beyond the stimulation artifact.

**Table 6:**
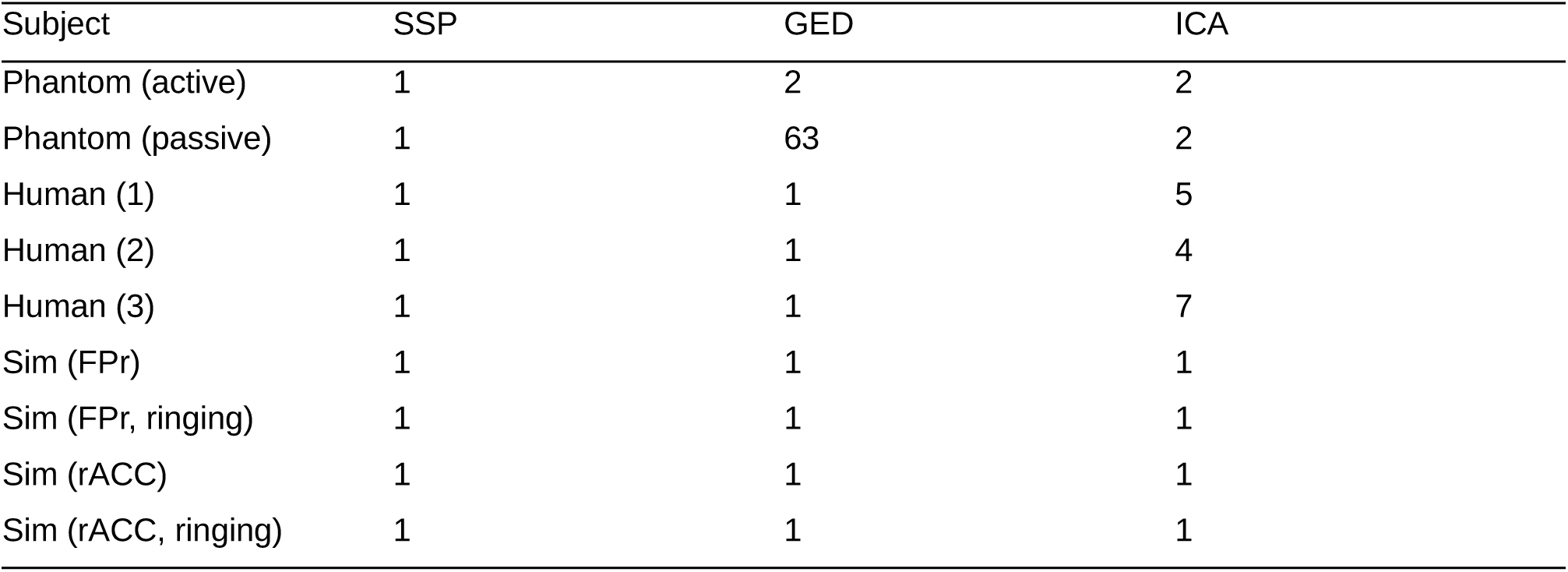
Number of Removed Artifact Components.

| Subject | SSP | GED | ICA |
| --- | --- | --- | --- |
| Phantom (active) | 1 | 2 | 2 |
| Phantom (passive) | 1 | 63 | 2 |
| Human (1) | 1 | 1 | 5 |
| Human (2) | 1 | 1 | 4 |
| Human (3) | 1 | 1 | 7 |
| Sim (FPr) | 1 | 1 | 1 |
| Sim (FPr, ringing) | 1 | 1 | 1 |
| Sim (rACC) | 1 | 1 | 1 |
| Sim (rACC, ringing) | 1 | 1 | 1 |

## Discussion

All methods were able to attenuate stimulation artifacts. However, notch filtering and spectrum interpolation, showed critical shortcomings, namely edge artifacts at the beginning and end of stimulation trains and attenuation of oscillatory activity spectrally overlapping with the stimulation. As more stimulation frequencies are beginning to be explored in taVNS, this detrimental effect on oscillations overlapping with brain activity is problematic.

Linear and mean interpolation are only applicable in selected circumstances, due to their destructive effects on the data. Both methods scale unfavorably at higher stimulation frequencies, requiring even more data to be removed.

AAS has proven to be useful, but has occasionally introduced residual or additional artifacts visible in the time domain, and in time-frequency representations. This may be due to imprecise detection of the stimulation onset or variability in the sampled artifact waveform. The need for an even more elaborate measurement setup, or manual fine-tuning of parameters, hinders the translation of this technique to less-than-ideal settings, such as clinical environments.

Zapline showed a loss of spectral power at higher frequencies, possibly due to the iterative application, with good recovery of oscillatory activity in phantom and simulated data. Power loss was evident at frequencies above 50 Hz. However, for many EEG analyses, the frequency range up to 40Hz is sufficient. An analytical advantage of the Zapline method is the preservation of the full data rank, which facilitates the later application of multivariate analysis. This comes at the cost of computational time, and the inability to implement this method in a real-time compatible manner.

The ICA noise removal resulted in a high similarity of the cleaned spectrum to the baseline across all datasets. However, manual component selection was required, and the artifact could appear in multiple components.

SSP showed strong artifact reduction across spectral and temporal domains, with minimal loss of data rank. However, as a blind separation method, it comes at the cost of possible aliasing of other signals, and enforces an orthogonality constraint on signal and noise subspaces that may not necessarily hold in real data.

Among all the methods, GED (SASS) has clear advantages. It has shown a recovery clean EEG in both the spectral and temporal domain. Since it involves the predefined comparison of artifact-containing and artifact-free covariance matrices, it is much less likely to remove non-artifact components compared to SSP or ICA. The use of permutation testing provides a simple and robust automated approach for estimating the number of artifact components. Unlike ICA, no manual selection of components is required, allowing for automated and more objective analyses. Similar to SSP, it is real-time compatible, since the necessary projection matrix for data cleaning can be estimated prior to an actual experiment and then applied in real-time to streamed data.

One limitation is that some methods, such as AAS and linear interpolation, could have been improved with manual fine-tuning. Similarly, the Zapline method could show some of its shortcomings in the higher frequency range due to the necessary iterative application. In the case of linear interpolation, fine-tuning the interpolation points may have resulted in a better representation of the underlying trends.

In conclusion, future studies using taVNS in combination with EEG, and possibly using real-time data streaming for stimulation parameters adaptation, should consider using GED (SASS) or SSP for data cleaning. Zapline could be considered as an alternative.

## CRediT authorship contribution statement

Joshua Woller: Conceptualization and design, Methodology, Investigation, Software, Formal analysis, Writing – original draft.

David Menrath: Conceptualization and design, Investigation, Writing – review & editing. Alireza Gharabaghi: Funding acquisition, Conceptualization and design, Project administration and supervision, Writing – review & editing.

This manuscript underwent meticulous revision by all authors and was further refined with regard to grammatical accuracy, sentence structure and wording using a convolutional neural network (DeepL) and an advanced language model (ChatGPT).

## Data and code availability

Data and code supporting this study, as well as custom implementations of the denoising algorithms, will be made available by the first author upon journal publication.

## Acknowledgements

The manuscript is based on work for the master thesis of J.W. at the University of Tübingen. We acknowledge support by the Open Access Publishing Fund of the University of Tübingen.

## Appendix A Thresholding Procedure & Segmentation

### Semi-Automated Annotation and Segmentation

A method was developed to infer both the onset of individual stimulation pulses and the beginning and end of stimulation trains based on the recorded EEG data. This segmentation into stimulation and non-stimulation periods is crucial for certain imputation methods such as linear interpolation or average artifact subtraction, but can also prove useful for SSP, GED and ICA.

One approach is to apply an additional artifact reference recording channel near the stimulation location, e.g. at the ear, to accurately sample the artifact signal and use that for later segmentation. However, the application of an additional channel takes time during the preperation. Further, it introduces a single, but not unlikely point of failure for analyses that rely on exact timing of stimulation pulses. Since ear anatomy differs between subjects, it can sometimes be difficult to find a location in which such a reference sensor can be attached practically, especially for smaller auriculae.

#### Generation of a synthetic artifact reference

A PCA-based approach was developed to create a synthetic reference channel for segmentation and annotation based on existing EEG data. While the stimulation artifact is clearly visible and stronger than the genuine EEG activity in *most* ipsilateral channels, it can be masked by other artifacts, such as muscle artifacts or electrode pop in individual channels, possibly hindering reliable detection by any single one channel. The procedure to generate a reliable artifact channel from the recorded EEG data is described in the following.

From the recorded data, a minimum of 3 channels (with a recommended amount of at least 5) that strongly show the stimulation artifact are chosen. An indicator for a suitable candidate is the clear presence of the rather weak impedance measurement pulses in the voltage trace. On data from these channels, a principal components analysis is performed. The section of the signal across which the PCA is computed may be specified so that the matrix is computed on an artifact epoch only, so that the PCA is fitted on the data from the stimulation epoch and applied to the entire data, yielding the component time courses.

For each of the resulting component time series, the power spectral density was computed using Welch’s method. Components are sorted according to the summed total signal at the stimulation frequency and its harmonics. The three components with the highest signal at those frequencies, as well as the average time course of those components were then saved. One of these components is chosen by the data analyst to be saved as an additional artifact reference channel in the data matrix of the EEG recording for later segmentation.

Usually, the first component provided a very good approximation of the artifact time course, with minor contributions from other noise sources. This procedure resulted in useable time courses of the artifact for simulated, human and phantom data, abolishing the need for an extra reference sensor to capture the stimulation artifact.

#### Peak Finding

The data from the artifact reference channel was then subjected to an automated iterative peak finding procedure to extract stimulation onset times precisely. Extracted peaks were then inspected for their conformity with known stimulation parameters.

The approach used an ascending threshold for peak heights, starting at 0.5 SDs above the channel mean and extending up to 1.9 SDs in steps of 0.2.

For each threshold (passed as the *height* parameter), peaks in the data were then extracted using scipy’s inbuilt peak finding algorithm (*scipy.signal.find_peaks)*, and the expected minimal horizontal distance between adjacent stimulation peaks, based on the stimulation frequency, was passed as the additional parameter *distance* to the function.

Each such detected peak was assigned one of two categories (stimulation pulse or impedance pulse) based on the distance to its neighboring peaks, which should be 0.04s for stimulation pulses at 25 Hz, and 1s for impedance measurement pulses.

These differences between peak times were also used to segment the data into stimulation and non stimulation segments. A peak time was considered the start of a stimulation train if its right neighbour peak occurred ∼1/25s later, and its left neighbour had more than twice that distance. This logic was inverted for the end of a stimulation train, i.e. the left neighbour has occurred ∼1/25s earlier, and the right neighbour was more than twice that distance away. The factor of 2 here was chosen heuristically.

For each iteration, the average distance among peaks of each class was calculated, and it’s absolute deviation from the expected value computed and stored as an error. Errors for both classes were added together. Additionally, non-detection of any class was penalized heavily, as was the failure of a threshold to induce a stable segmentation into stimulation periods as defined above (i.e., a threshold resulting in an unequal amount of onsets and ends of stimulation trains). For each off these, an additional error of 10,000 was added for this iteration, effectively preventing such segmentation from being chosen later.

After all iterations ran through, the threshold with the minimal error was chosen for the final segmentation. The peak times were written to stimulus channels and appended to the data, leading to one stimulus channel indicating the occurrence of any peak (impedance and stimulation), and separate channels for each pulse class as well. Additionally, detected onsets and ends of stimulation trains were used for annotations of data segments as stimulation periods using mne-Python.

The data annotated in such manner could be then passed to the different denoising procedures.

## Appendix B

### Data Simulation

#### Simulation of taVNS artifacts in EEG data

For each 5 seconds stimulation train, equidistant stimulation onsets for individual pulses were generated, resulting in an impulse train which was then convolved with a biphasic kernel of unit amplitude and 2ms duration. This duration deviates from the pulse duration of the device as the artifact is in fact extended in time in the recording (4-5ms artifact duration in the recording compared to 250μs pulse duration of the device). After each stimulation segment, a 5s pause segment was appended. In these non-stimulation segments, impedance measurement pulses at 1Hz were simulated according to the above procedure, but at 1/3 of the unit amplitude. The resulting artifact time course was resampled at 1kHz using an antialiasing filter. This induced some desired smoothing and ringing of the artifacts.

#### Construction of the simulated EEG signal

The artifact time course was first scaled to be within the amplitude range of EEG data, so that the artifact had a maximal amplitude range of 160µV. Then for each sensor location, its relative distance in the flattened x-y plane of sensor locations relative to a reference sensor with maximal artifact amplitude (chosen arbitrarily, here “EEG 018”) was calculated, and used to generate weights from 1 to 0 with which the artifact time course was then multiplied and added to the sensor data.

For each simulated source, two simulation versions were produced. One with full ringing, and one in which the ringing was reduced by applying a gaussian envelope of 15ms FWHM around each stimulation pulse onset after the resampling and scaling of the data, but before adding it to the sensor signals.

## Appendix C

### Modified Artifact Subtraction

For average artifact subtraction, it is necessary to epoch data around individual stimulation pulses. This can however lead to additional problems. As the stimulation frequency lies in the same range as brain oscillatory activity, such as the beta range and the classic 25Hz stimulation frequency, sampling the artifacts might also sample underlying synchronised oscillatory neural activity. This problem is only exacerbated if one considers such activity to be evoked or perhaps the result of entrainment to the stimulation frequency. Blindly applying AAS in this context might introduce additional artifacts at best, and systematically affect the interpretation of the findings at worst. To counteract this, a modified approach was developed. Additionally to the artifact epochs outlined above, non-stimulation epochs of the same duration as artifact epochs are created. These are sampled in the gaps between the regularly spaced stimulation artifacts. From these epochs, a correction template for the artifact template is derived. If there is no entrained oscillatory neural activity, this correction template will only contain minor noise contributions. If there however is underlying oscillatory activity phase-locked to the stimulation, this template will lead to an average signal that is approximately the same as the aliased activity in the artifact template, up to the sign. This sign uncertainty is due to the fact that these two epoch types may sample either the same phase of a neural oscillation, e.g. if the oscillatory activity is an even-numbered harmonic of the stimulation frequency, or in anti-phase, if it is an odd-numbered harmonic. The sign of the dot-product between both average vectors can then be used to decide whether the correction template is added (negative sign) or subtracted (positive sign) from the artifact template.

Specifically, this modified approach deals with the problem of a template aliasing neural signal for ongoing symmetric oscillatory activity. It does not solve however the problem of evoked activity being averaged together with the stimulation artifact, which is an inherent limitation of artifact template subtraction approaches.

